# Humans are resource-rational predictors in a sequence learning task

**DOI:** 10.1101/2024.10.21.619537

**Authors:** Vanessa Ferdinand, Amy Yu, Sarah Marzen

**Author notes:** Contributing authors.

## Abstract

Organisms can solve complex tasks despite having limited cognitive resources when those resources are used optimally. Doing so optimally makes an organism “resource-rational”. In this paper, we show for the first time that humans are resource-rational at prediction. In a novel sequence learning experiment, participants predict data generated from hidden Markov models (HMMs) and receive online feedback via clicker training. We compute the predictive rate-accuracy (PRA) curve for each HMM to solve for the highest accuracy achievable for a given cognitive capacity, or “rate”, quantified as the mutual information between participants’ predictions and the underlying causal state of the sequence being predicted. We found that the majority of participants achieve near-optimal prediction with various cognitive capacities, despite performing well below the maximum predictive accuracy on the task overall. We also show that this information-theoretic approach to measuring cognitive capacity can be grounded in the established psychological science concept of working memory: participants who extracted higher quantities of mutual information in the sequence learning task showed significantly higher working memory in a complex digit span test. This research provides new avenues for assessing smart behavior in difficult prediction tasks, provides a new methodology for assessing resource-rationality, and provides evidence that humans are resource-rational predictors.

## 1 Introduction

A surprising amount of human behavior relies on prediction. Some of this prediction occurs unbeknownst to us in algorithms that we commonly use, such as recommender systems, weather applications, and financial management tools. But most of this prediction is done by us ourselves, nearly implicitly, such as predicting the movement of vehicles on the road, the social behavior of other humans, or the structure of linguistic utterances. There are strong theoretical reasons for why prediction would be so important to organisms. For instance, humans are thought to be reinforcement learners [1] who survive and procreate by maximizing the reward accrued over their lifetime, and reinforcement learning algorithms rely on either implicit or explicit predictions of the world [2].

Not all information about the past is useful for predicting the future [3]. Since organisms have limited capacities for processing information [4, 5], they must work out what information is worth encoding, and what is not, in order to make efficient predictions about the future. This idea is an information-theoretic instantiation of resource rationality [6, 7] or computational rationality [8, 9]. Using information-theoretic quantities as perceptual costs has allowed researchers to explain a number of empirical findings in a wide variety of areas, including various aspects of macroeconomic behavior [10, 11], Shepard’s universal law of generalization [12], human decision making in simple choice tasks [13], the fuzziness of color naming systems [14], and a number of empirical findings on neural coding and working memory [15]. And, while not done on humans, recent work has shown that salamander retinal ganglion cells [16] and cultured cortical neurons from rats [17] both predict stimuli efficiently in an information-theoretic sense. Even bacteria appear to nearly maximize their performance in a chemotactic gradient-climbing task given limitations on the rate of information they take in about the environment [18]. With regards to prediction, information is a resource that must not be used lightly, but instead must be allocated properly so that all information about the past is used to effectively predict the future.

To study memory and prediction, we treat prediction as a forecasting of time series data and investigate this through sequence learning experiments in which the human or a human-crafted algorithm is shown some number of symbols or numbers and asked to predict the next batch of symbols. These symbols are generated from special types of hidden Markov models (HMMs), which generate infinite-order Markov processes, unlike prior seminal work which focused on Markovian stimuli [19]. The infinite-order Markov stimuli can be as simple or complex, as artificial or naturalistic, as one wishes, simply by changing the topology and transition probabilities of the underlying hidden Markov model. This allows us to show that humans are sensitive not just to transition probabilities of hidden Markov models [19], but also to constantly changing predictive features of the input encapsulated by the hidden states themselves.

In this paper, we use a novel sequence learning paradigm with infinite-order Markov stimuli to test whether human learners are able to extract information for predicting timeseries data in a resource-rational way. We used HMMs to generate long sequences of observable outcomes from hidden causal states. All participants gradually learned to predict the observable outcomes over time, but with varying degrees of success. We used each participant’s observed prediction behavior to measure the amount of causal information they extracted by calculating the mutual information between their predictions and the HMM’s causal states. To assess the resource-rationality of their behavior, we applied a new computational method [20] that extends the information bottleneck method [21] to predictive information extraction. For each demonstrated information processing capacity, this method solves the maximum predictive accuracy achievable given that capacity and the HMM in question. The result is a *predictive rate-accuracy* (PRA) curve that defines the range of optimal and achievable solutions over all possible information processing capacities, running between maximization behavior (as the best strategy that carries zero predictive information about the causal states in a timeseries) and the “ideal strategy” (which achieves the highest accuracy possible for the HMM by using the largest information processing capacity necessary). Rather than deem participants to be poor predictors for not discovering the ideal strategy, we instead asked: how close is each participant’s performance to optimal for their demonstrated level of predictive information extraction? Evaluating a learner’s performance in this way can help researchers identify the smart strategies that resource-limited organisms use and distinguish them from behavior that is truly sub-optimal for the task at hand.

## 2 Results

### Participants learned to predict each timeseries

We found that participants were able to predict the sequences generated from all three HMMs (Fig. 1, top row), but with varying degrees of success. Participants’ average predictive accuracy was 88% for NP, 66% for DP, and 58% for EP. For comparison, the ceiling on predictive accuracy in each condition is 95% for NP, 81.25% for DP, and 76.92% for EP: these are the scores participants could have achieved if they were using the ideal strategy to predict each HMM.

**Fig 1.**
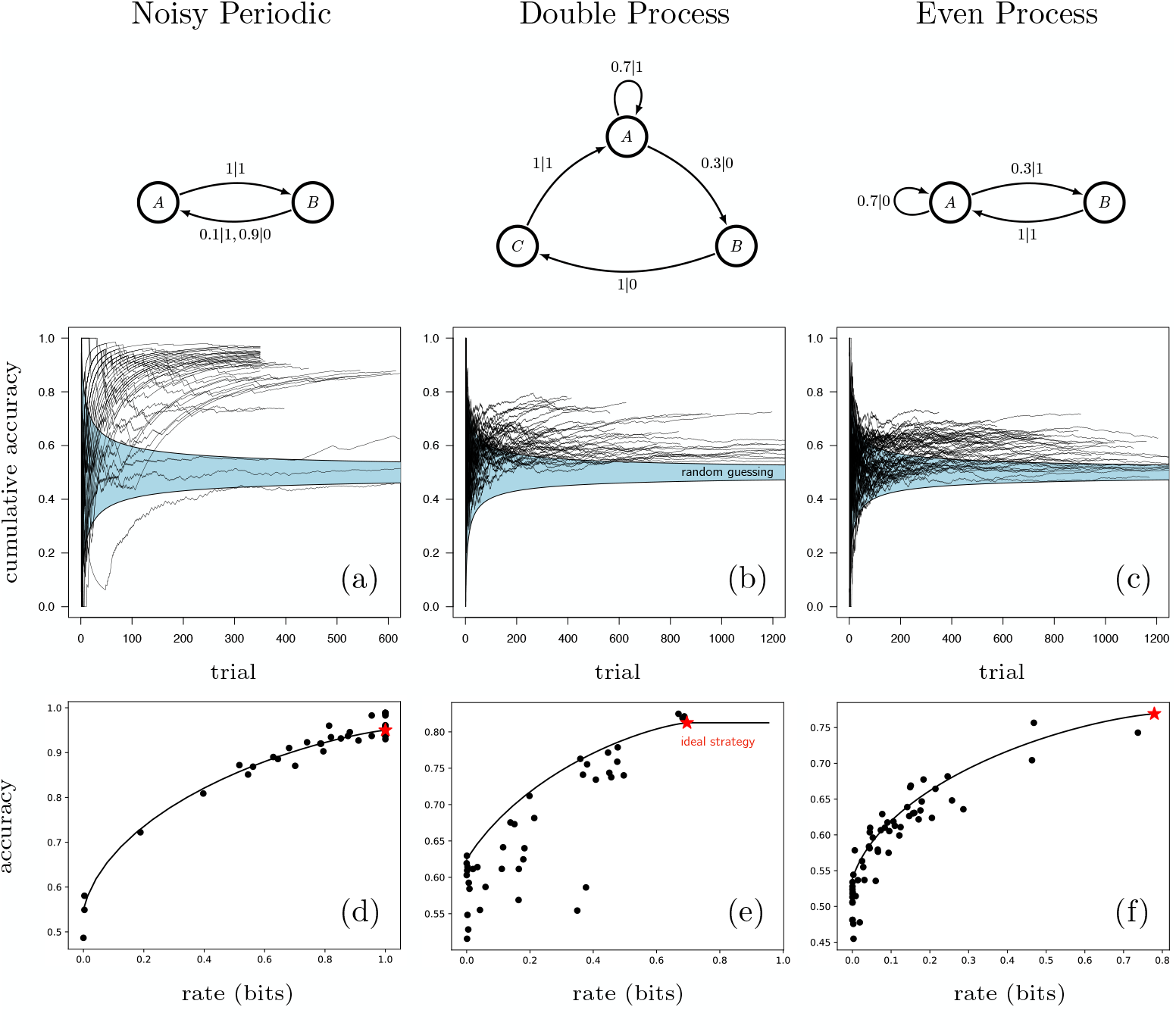
*Top row:* In each experimental condition, participants were trained on sequences generated by one of these three hidden Markov models: the Noisy Periodic Process (NP), the Double Process (DP) and the Even Process (EP). *Middle row:* Cumulative accuracy scores by trial for sequences in condition NP (a), DP (b), and EP (c). Each black line shows the learning trajectory of one participant. Over time, nearly all participants learned to predict the sequence above chance levels (blue region is the 95% confidence interval around random answering). The predictive accuracy ceiling for each process is a) 95%, b) 81.25%, and c) 76.92%. *Bottom row:* Predictive rate-accuracy curves along with points showing participant performance in each condition. Participants perform optimally (i.e. close to the curve) in NP (d) and EP (f) but exhibit sub-optimality in DP (e). In NP, participants obtain high accuracy with memory-intensive strategies, whereas in EP they tend to achieve lower accuracy with relatively memory-less strategies.

Fig. 1a-c shows each participant’s cumulative accuracy over time, where each black line is the learning trajectory of one participant. The blue zone shows the region of cumulative accuracy scores that would be expected if participants were randomly choosing between the two responses with equal probability (i.e. the binomial 95% confidence interval given *N* trials). Across all conditions, 127 of the 140 trajectories eventually exited the blue zone, meaning these participants learned to predict the HMMs. In NP, learning trajectories reached higher accuracy levels faster, indicating this was the easiest timeseries for participants to predict. In EP, trajectories were slowest to exit the blue zone and reached lower maximum accuracy levels, indicating this was the hardest timeseries to predict.

### Participants used resource-rational prediction strategies

Participants discovered a variety of optimal prediction strategies over a range of information-processing capacities. Figure 1d-f shows how participants performed relative to the PRA curve for each HMM. Each black dot shows the computed rate and accuracy for the last half of each participants’ prediction trials.

To understand how close to optimal participants’ predictions were in each condition, we computed their relative distance from the curve along the accuracy dimension (the y-axis in Fig. 1d-f) [22]. Fig. 2a shows these distances from the curve per condition. Mean distance from the curve was 0.00 for NP, 0.07 for DP, and 0.03 for EP. The results of a linear mixed effects model show that predictions in the NP condition was not significantly different from optimal and that predictions in DP and EP were close to optimal, but significantly below the curve. The estimate for distance from the curve was -0.003 for NP (*SE* = .007, *t* = −0.392, *p* = 0.695), 0.066 for DP (*SE* = 0.008, *t* = 8.448, *p <* .001), and 0.028 for EP (*SE* = 0.007, *t* = 4.328, *p <* .001). Distance from optimal also differed significantly between the three conditions. EP was significantly less optimal than NP (Estimate = 0.031, *SE* = 0.010, *t* = 3.184, *p* = .002), DP was significantly less optimal than NP (Estimate = 0.069, *SE* = 0.011, *t* = 6.405, *p <* .001), and DP was significantly less optimal than EP (Estimate = 0.037, *SE* = 0.010, *t* = 3.667, *p <* .001). Taken together, these results mean that participant’s predictions in NP were optimal, predictions in EP were less optimal, and predictions in DP were even less optimal with an effect size 2.4 times worse than EP. Optimality rankings between conditions is summarized as NP *>* EP *>* DP.

**Fig 2.**
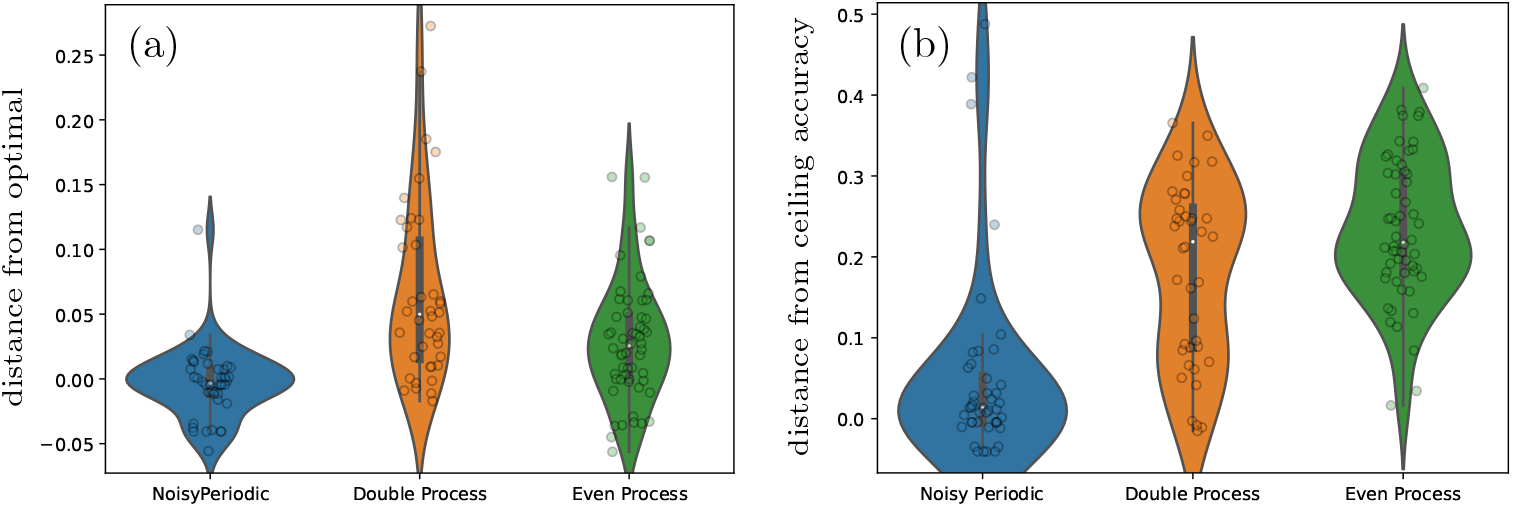
(a) Optimality of participants’ prediction behavior measured as distance from the PRA curve in each condition. Distances have been normalized across curves to be directly comparable between conditions. Predictions for NP are closest to optimal and those for the DP are farthest from optimal. (b) A traditional analysis of performance on each sequence type. The y-axis shows how far participants’ predictions were from the maximum achievable predictive accuracy per condition, without taking each participants’ rate into account. Each condition elicited significantly different performance. Prediction in NP is significantly better than in DP and EP, but no significant difference is detected between DP and EP.

### A traditional analysis of prediction performance

To highlight what the resource-rational analysis buys us, we compare our PRA results to those of a traditional analysis of performance in sequence learning tasks, where participant performance is simply compared to the accuracy ceiling in each condition *without* accounting for each participant’s unique rate.

Fig. 2b shows the distances between participants’ accuracy scores and the accuracy ceiling for each HMM (i.e. the accuracy under the ideal strategy for predicting the HMM). We re-ran the linear mixed effects analysis with this distance as the dependent variable and found significant differences between all conditions except DP and EP. The estimate on distance from ceiling was 5% for NP (*SE* = 0.014, *t* = 4.041, *p <* .001), 15% for DP (*SE* = 0.014, *t* = 10.493, *p <* .001), and 18% for EP (*SE* = 0.012, *t* = 14.901, *p <* .001). Distance from ceiling differed significantly between the three conditions. EP was significantly worse than NP (Estimate = 12.6%, *SE* = 0.018, *t* = 7.203, *p <* .001), DP was significantly worse than NP (Estimate = 9.3%, *SE* = 0.019, *t* = 4.879, *p <* .001), and EP was marginally worse than DP (Estimate = 3.3%, *SE* = 0.018, *t* = −1.815, *p* = 0.07) but for the EP-DP comparison, the difference is not significant. Going off of the estimates alone, the traditional analysis re-ranks participant performance as NP *>* DP *>* EP, but in terms of statistical significance it does not find a difference between EP and DP, thus NP *>* DP = EP. In short, the traditional analysis fails to pick up on the efficient but low-accuracy strategies used by participants in EP and classifies them all as poor performance.

### Occasional discovery of the ideal strategy

The red stars in Fig. 1d-f correspond to the ideal prediction strategy for each HMM. The ideal strategy for any HMM is solved as the strategy that produces the maximum-likely symbol given each hidden state (i.e. state-dependent maximization behavior). Several participants discovered the ideal strategy over the course of their session, although not all participants stuck with it. In NP, 69% of participants produced runs of the ideal strategy more than expected by chance and 56% of them stuck with it for at least 50 trials. In DP, these numbers were 85% of participants at 31% persistence. In EP, they were 66% of participants at 9% persistence. All of the best-performing participants (in the top right corner on the PRA curves) used the ideal strategy for the majority of their trials. See Supplementary Information for a detailed discussion of nonstationarity (i.e. strategy-switching across trials), its size in this experiment, and its effects on assessments of optimality.

### Working memory is related to rate

We investigated the relationship between rate (an information-theoretic quantity related to memory capacity which was computed directly from participants’ prediction behavior in the sequence learning experiment) and an independent, standardized measure of participants’ working memory capacity developed by psychologists: the complex digit span test. The results of a linear mixed effects model and a likelihood ratio test showed that working memory scores were significantly related to rate (*χ*^2^(1) = 4.329, *p* = 0.037). The effect size shows a one point increase in working memory score leading to an estimated 0.03±0.014 (standard errors) point increase in rate. This is equivalent to a 2.4% increase in score yielding a 3% increase in rate. This result means that our method calculating participants’ rate from their observed prediction behavior is, indeed, an indicator of participants’ information processing capacity and can be grounded in the established psychological sciences concept of working memory capacity.

### Prediction algorithm results

Resource-rationality is a computational-level framework: it can tell us about the optimality of behaviors without telling us, or even requiring that we know, what is happening at the algorithmic-level (see Marr’s levels of analysis for cognitive systems [23]). This means that a separate analysis is required to understand which strategies participants were using to acheive resource-rational prediction. We tested four classes of prediction algorithms: Bayesian order-R Markov modeling (n-gram strategies), Bayesian Structural Inference (BSI), Long Short-Term Memory Units (LSTMS), and logistic regression.

Fig. 3 shows the best-fit model per participant. Among the four algorithms we tested, we found that participants were most likely to have used the n-gram algorithm, and by a wide margin as well. Typically, the best-fit model for each participant greatly dominated in terms of likelihood and thus probability, as measured by Δ*AIC* and *wAIC* [24], with most Δ*AIC* values for the second-best model falling far above Δ*AIC* = 4. Two models are considered equally likely when Δ*AIC* scores are less than 2 and only three participants fell into this range. For these three cases, we report 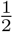 participant per best-fit and second-best-fit model in Fig. 3. Interestingly, we typically found that participants did not drop data or bias themselves towards simpler models when using n-gram or BSI strategies.

**Fig 3.**
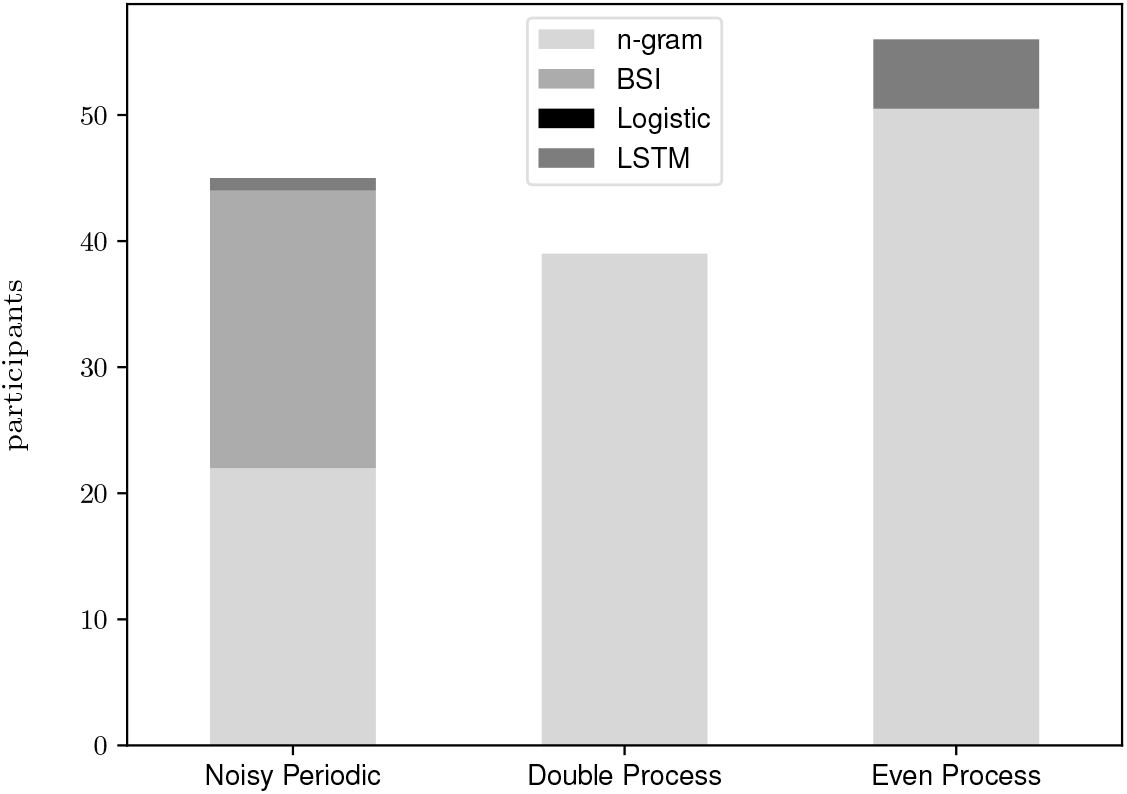
The number of participants, by experimental condition, whose prediction behavior was best-described each of the four prediction algorithms investigated: Bayesian order-R Markov modeling (n-gram), Bayesian Structural Inference (BSI), Long Short-Term Memory Units (LSTMS), and logistic regression (Logistic). Participants’ prediction strategies were most consistent with *n*-gram strategies.

## 3 Methods

All experimental protocols were approved by the University of Melbourne Human Research Ethics Committee (1954586.1) and the United States Air Force Human Research Protections Office (FWR20190129X). The methods were carried out in accordance with Australia’s National Statement on Ethical Conduct in Human Research 2007 and its subsequent revisions. Informed consent was obtained from all participants.

### Sequence generation

Participants were tasked with predicting sequences that were generated by hidden Markov models (HMMs) [25]. Such models allow for generation of sequential stimuli that quantifiably vary from low complexity to high complexity, from low randomness to high randomness, and from low predictability to high predictability [26]. Furthermore, generation of these stimuli is computationally efficient and does not require the experimentalist to cleverly hand-design certain patterns: the patterns emerge from the structure of the model itself, which is easily visualized and captured in the diagrams shown in Fig. 1 (top row).

Each participant received a single long sequence from one of the three HMMs generated in the following manner: a hidden state (*σ*) was randomly selected and an emitted symbol (*x*) was drawn with probabilities given by *p*(x|σ) as shown on the arrows in the HMM diagrams. The next hidden state was transitioned to according to *p*(*σ*_*t*+1_|*x*_*t*_, *σ*_*t*_) and the process repeated 3000 times, creating a sequence of length 3000. Participants were trained on the sequence of emitted symbols only and had no access to the hidden states. Table 1 shows an example sequence of length 12 generated from the Even Process and the relationship between the hidden state, emitted symbol, and a participant’s prediction. In this example, the participant makes correct predictions on all trials except *t* = 3, 5 and 10. The prediction sequence shown is the ideal strategy for the Even Process and can be verbalized as “win-stay, lose shift-once”. Here, we see that the ideal strategy perfectly the tracks the hidden states, but not the emitted symbols.

**Table 1.**
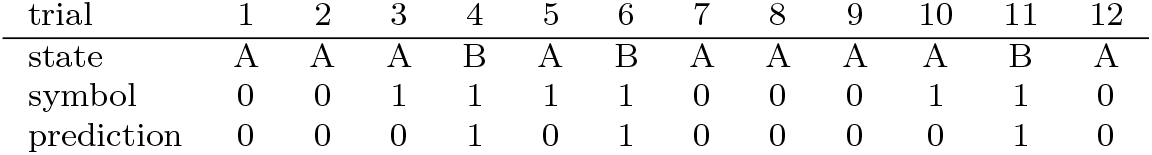
Example Even Process sequence of hidden states, emitted symbols, and a participant’s predicted symbols.

### Sequence learning experiment

93 participants took part in the experiment and were randomly assigned to one of three conditions corresponding to one of the three HMMs in Fig. 1 (top row). Participants were allowed to take the experiment as many times as they wanted, resulting in 140 completed sessions: 44 in the NP condition, 40 in DP, and 56 in EP. Participants were recruited using Amazon’s crowdsourcing platform MTurk and paid above Federal Minimum wage for their time. The experiment was administered over a minimalistic computer interface where participants made predictions by pressing one of two buttons on their keyboard (F and J) and viewed a dial on the screen showing the cumulative accuracy of their predictions. Participants were instructed to “try to discover the pattern” and began by pressing any button (F or J). The timing of each button press was determined by the participant and corresponded to one trial. The assigned HMM determined which of the two buttons was “correct” on any given trial. For example, if the *i*th emission from the HMM was symbol 1, then the correct key on the *i*th trial was key 1 (counterbalanced between participants to F or J). If the participant pressed the correct key, they would hear a clicker sound. Otherwise, they would hear nothing. No emission symbols were shown on the screen; these symbols only determined which key was correct on each trial. We used a self-paced design and allowed participants to complete as many trials as they wanted, capped at 2700 trials. If participants reached a certain cumulative accuracy threshold (80% for NP and 70% for DP and EP), they received an additional 1 USD reward and the experiment automatically ended after 300 more trials were collected. The number of trials the participants completed per condition were NP: mean = 494 (min = 123, max = 2700), DP: mean = 914 (min 323, max = 2700), EP: mean 907, (min = 56, max = 2700).

### Working memory test

Individual differences in participants’ working memory were assessed with a complex digit span test [27, 28]. 37 participants opted in to taking the working memory test as well as the sequence learning experiment. Participants were presented with a series of digits and asked to recall them in order. Between each presentation and recall phase of a trial, a distractor task occurred where participants were shown a sentence and evaluated whether it was grammatical or not. The number of digits to be recalled per trial started at 2 and gradually increased to 5, totalling 42 digits across all trials. The working memory score is simply the number of digits, out of 42, that the participant accurately recalled.

### Predictive rate-accuracy curves

Given a certain amount of cognitive resources, are humans predicting as well as possible? We quantify cognitive resources using a measure of memory previously used for salamander retinal ganglion cells [16] and cultured neurons [17]– the mutual information between the state of the sensor, proxied by the human’s prediction, and the forward-time causal state of the input sequence [29]. The forward-time causal state is the minimal sufficient statistic for prediction of the input sequence [29], and so the nonlinear correlation [30] between the human’s behavior and this minimal sufficient statistic of prediction is an information-theoretic rate that can quantify memory. We quantify predictive power simply by the predictive accuracy. For any given rate, there is a maximal predictive accuracy attainable. We calculate this maximal predictive accuracy and show human behavior relative to the resultant predictive rate-accuracy (PRA) curve [22]. The input sequences are specially designed so that this calculation can be performed [20]. Plotted next to the PRA curve are the actual rates and accuracies of humans; if the humans are close to the curve, then they are resource-rational or efficient predictors. Because we collected a finite set of data from each participant, points can fall above the curve due to chance fluctuations in accuracy scores.

### Prediction algorithms

A maximum likelihood method [31] adjusted to become a maximum a posteriori method was used to infer which prediction strategy was being employed by participants. There were four candidates: Bayesian order-*R* Markov modeling [32] or n-gram observers, Bayesian Structural Inference [33], Long Short-Term Memory Units [34], or logistic regression. For all classes of algorithm considered, we assume the participant trains the model on the history of symbols emitted across all previous trials and then uses that model to predict the next trial. We report the “best-fit” model as the one (from this set of four) that minimizes the Akaike Information Criterion (AIC), which is 2*K* − 2 log *L*, where *K* is the number of parameters and *L* is the probability of the data given the model [24].

## 4 Discussion

This research showed, for the first time, that humans learn to predict time-series data using a range of optimal resource-rational strategies. Analyzing prediction behavior in a resource-rational framework provided new insight into how difficult various stochastic processes are for humans to learn. The three Hidden Markov Models (HMMs) we chose to test participants on varied in difficulty, in the sense that they each had different ceilings on predictive accuracy. Based on these values, DP should have been easier to predict than EP, but we found that participants discovered significantly more resource-rational strategies for EP than for DP, indicating that the structure in EP is easier for participants to learn. Furthermore, we found that a traditional analysis of prediction success, measured by how close participants get to the accuracy ceiling for the task, showed no difference between DP and EP and would incorrectly deem these two tasks equally hard for participants to learn. Not only would an analysis using absolute prediction fail to pick up on the resource-rational prediction of the participants, but also it would have ranked the tasks in terms of their difficulty for participants differently.

Resource-rational prediction occurs when learners achieve the highest predictive accuracies possible under a given cognitive capacity for collecting, processing, or storing information [6]. However, it can be challenging to quantify an organism’s cognitive capacity and calculate its associated predictive performance optimum. We operationalized the notion of cognitive capacity as a *rate* [20] given by the mutual information between the causal states in a sequence-generating HMM and the participant’s predictions of that sequence. This rate quantity has been used in prior work to establish the resource-rational prediction capabilities of salamander retinas [16] and cultured neurons [17] but had not yet been applied to prediction behavior at the level of entire organisms. We also showed that rate correlated with individual differences in working memory: participants who extracted higher quantities of mutual information in the sequence learning task showed significantly higher working memory in a complex digit span test. This finding grounds the information-theoretic quantity, rate, in the established psychological science concept of working memory and validates the use of this metric as proxy for cognitive capacity. As for our chosen measure of predictive success, we simply used predictive accuracy as a metric for how well a participant predicted, though other informational measures could have been used instead [14, 16, 17]. With these substitutions, testing resource-rational prediction becomes an extension of the well-worn field of rate-distortion theory [20], meaning ideas from the information theory subfield of lossy compression [35] can be used to understand human cognition. One simply measures the closeness of participant data to the predictive rate-accuracy curve to assess whether or not those participants are resource-rational predictors. It is an open question why quantities that normally apply in the asymptotically blocklength limit should apply to the single-symbol code used by humans, though at the very least rate is a nonlinear correlation coefficient [30] that– from our analysis– captures memory.

There is a large body of research on sequence learning in the fields of implicit learning [36], statistical learning [19, 37, 38], and artificial grammar learning [39]. However, the resource-rationality angle is just not present in this body of research. As such, the present study represents a departure from this literature that accords with new ideas from both economics [10] and biology [12, 14–18]. A resource-rational framework could easily be imported into existing sequence learning methodologies, especially because so many of them use finite-state automata [39–44] and HHMs [45] to generate their sequence learning stimuli. Such studies typically focus on how close participants come to learning the true target grammar that generated the sequence, and they do so by analyzing reaction times [41], testing learners’ sensitivity to transition probabilities [19, 46], or fitting an HMM to participant data and assessing its distance from the target HMM [45]. Our framework can complement these methodologies by providing a way to assess the optimality of intermediate strategies as they approach the target strategy or stable strategies that never reach the target. The vast majority of sequence learning paradigms focus on Markovian stimuli [46]. While not as well-understood, HMM-generated stimuli are more complex than Markovian stimuli and offer many more points of contact with topics in memory, prediction, and how learners extract information about the casual states behind timeseries data [29].

Comparative cognition is one area where this framework could have substantial impact, as it allows one to locate the performance of diverse cognitive systems on a single landscape of optimal and sub-optimal prediction strategies. Just as we showed that individual humans used a variety of smart strategies across different cognitive capacities, future research could show how various species of non-human animals and artificial systems cluster in different areas of this space. Resource-rationality is even useful for engineered prediction devices, as they too have finite memory [47, 48] and the predictive performance of different machine learning algorithms can also be compared within this framework [22]. Comparing the performance of humans and machine learners in prediction tasks could be useful for the development of complementary human-computer interaction systems that leverage the unique abilities of both cognitive systems by assessing where in the PRA space humans and machine learners fall when predicting on their own, and where they fall when predicting in tandem. By the same rationale, this method could be extended to understand predictive performance in collective intelligence [49] and wisdom-of-crowds phenomena [50] simply by letting a group of interacting organisms make predictions.

In addition to the computational-level analyses discussed above, we separately addressed the question of what strategy was being used to achieve resource-rational prediction at an algorithmic level [23] by conducting a maximum likelihood analysis of four candidate prediction algorithms [31]. We found that participant behavior was most consistent with order-*R* Markov models [32] and that this was the best-fit strategy across all three experimental conditions. A few participants were fit better by state-of-the-art recurrent neural networks [34] or state-of-the-art Bayesian inference algorithms [33], but no participants were fit better by logistic regression strategies. This order-*R* Markov model result means that participants behaved like “n-gram learners” and looked for patterns in sub-strings of symbols to make their next guess. This finding is consistent with a large body of evidence from statistical learning experiments showing that human participants engage in “chunk learning” [51–54] and track superficial statistics in sequential stimuli that correspond to bigrams, trigrams, etc [42, 55].

In the future, the paradigm introduced here could be used with additional measurements and more complex stimuli to gain more insight into brain activity and cognition. Brain scans could be used to provide a better estimate of the cognitive resources participants use to complete the task. Sequence learning tasks with more salient and less artificial input could be used to understand how saliency and priors on input affect their ability to produce efficient predictions. More naturalistic input would likely lead to less nonstationarity in the prediction strategy by the end of the task and would enable better determination of resource-rationality, even though those prediction tasks would be harder, since organisms would already be familiar with the task in some way.

Our finding that humans are resource-rational predictors in artificial but complex sequence learning tasks is a useful step towards an understanding of Marr’s computational level, and our maximum likelihood method approach to understanding the participants’ prediction and learning strategies is a useful step towards an understanding of Marr’s algorithmic level [23]. It may be the case that humans are resource-rational decision makers and that simple controlled environments like the ones used here show if and how this is true.

## Data Availability

(to be released upon publication)

## Code Availability

(to be released upon publication)

## Acknowledgements

Special thanks to research assistant employees Jacob Kuek and Malinga Perera. JK recruited and managed participants, managed data collection, and programmed the working memory test. MP programmed the sequence learning experiment and deployed it on Google App Engine. S. Marzen acknowledges valuable conversations with Stephanie Palmer and Piper Connelly, and S. Marzen and A. Yu acknowledge valuable conversations with Mark Huber. This study was supported by the US Air Force Office for Scientific Research, Grant Number FA9550-19-1-0411. Open access funding was supported by the Melbourne School of Psychological Sciences.

## Author contributions

V.F. conceived of and designed the experiment, supervised research assistants to program and run the experiment, designed some of the analyses, analyzed the sequence and working memory data, and wrote and edited the paper. S.M. conceived of the experiment, designed some of the analyses, analyzed some of the sequence data, and wrote and edited the paper. A.Y. analyzed the sequence data and contributed expertise on training neural networks. All authors have contributed to, read, and approved the final version of the manuscript.

## Additional information

**Supplementary Information** is available online at (to be released upon publication).

**Correspondence** should be addressed to S.M.

## References

[1] Silver, D., Singh, S., Precup, D., Sutton, R.S.: Reward is enough. Artificial Intelligence 299, 103535 (2021)

[2] Sutton, R.S., Barto, A.G.: Reinforcement Learning: An Introduction. MIT Press, ??? (2018)

[3] Bialek, W., Nemenman, I., Tishby, N.: Predictability, complexity, and learning. Neural computation 13(11), 2409–2463 (2001)

[4] Barlow, H.B., et al.: Possible principles underlying the transformation of sensory messages. Sensory communication 1(01), 217–233 (1961)

[5] Shannon, C.E.: A mathematical theory of communication. The Bell system technical journal 27(3), 379–423 (1948)

[6] Lieder, F., Griffiths, T.L.: Resource-rational analysis: Understanding human cognition as the optimal use of limited computational resources. Behavioral and brain sciences 43, 1 (2020)

[7] Icard, T.F.: Resource rationality (2023)

[8] Lewis, R.L., Howes, A., Singh, S.: Computational rationality: Linking mechanism and behavior through bounded utility maximization. Topics in cognitive science 6(2), 279–311 (2014)

[9] Gershman, S.J., Horvitz, E.J., Tenenbaum, J.B.: Computational rationality: A converging paradigm for intelligence in brains, minds, and machines. Science 349(6245), 273–278 (2015)

[10] Sims, C.A.: Implications of rational inattention. Journal of monetary Economics 50(3), 665–690 (2003)

[11] Sims, C.A.: Rational inattention: Beyond the linear-quadratic case. American Economic Review 96(2), 158–163 (2006)

[12] Sims, C.R.: Efficient coding explains the universal law of generalization in human perception. Science 360(6389), 652–656 (2018)

[13] Callaway, F., Hardy, M., Griffiths, T.L.: Optimal nudging for cognitively bounded agents: A framework for modeling, predicting, and controlling the effects of choice architectures. Psychological Review (2023)

[14] Zaslavsky, N., Kemp, C., Regier, T., Tishby, N.: Efficient compression in color naming and its evolution. Proceedings of the National Academy of Sciences 115(31), 7937–7942 (2018)

[15] Jakob, A.M., Gershman, S.J.: Rate-distortion theory of neural coding and its implications for working memory. Elife 12, 79450 (2023)

[16] Palmer, S.E., Marre, O., Berry, M.J., Bialek, W.: Predictive information in a sensory population. Proceedings of the National Academy of Sciences 112(22), 6908–6913 (2015)

[17] Lamberti, M., Tripathi, S., Putten, M.J., Marzen, S., Feber, J.: Prediction in cultured cortical neural networks. PNAS nexus 2(6), 188 (2023)

[18] Mattingly, H., Kamino, K., Machta, B., Emonet, T.: Escherichia coli chemotaxis is information limited. Nature physics 17(12), 1426–1431 (2021)

[19] Saffran, J.R., Aslin, R.N., Newport, E.L.: Statistical learning by 8-month-old infants. Science 274(5294), 1926–1928 (1996)

[20] Marzen, S.E., Crutchfield, J.P.: Predictive rate-distortion for infinite-order markov processes. Journal of Statistical Physics 163, 1312–1338 (2016)

[21] Tishby, N., Pereira, F.C., Bialek, W.: The information bottleneck method. arXiv preprint physics/0004057 (2000)

[22] Marzen, S.E., Crutchfield, J.P.: Probabilistic deterministic finite automata and recurrent networks, revisited. Entropy 24(1), 90 (2022)

[23] Marr, D.: Vision: A Computational Investigation Into the Human Representation and Processing of Visual Information. MIT press, ??? (1982)

[24] Anderson, D., Burnham, K.: Model selection and multi-model inference. Second. NY: Springer-Verlag 63(2020), 10 (2004)

[25] Rabiner, L.R.: A tutorial on hidden markov models and selected applications in speech recognition. Proceedings of the IEEE 77(2), 257–286 (1989)

[26] Feldman, D.P., McTague, C.S., Crutchfield, J.P.: The organization of intrinsic computation: Complexity-entropy diagrams and the diversity of natural information processing. Chaos: An Interdisciplinary Journal of Nonlinear Science 18(4) (2008)

[27] Turner, M.L., Engle, R.W.: Is working memory capacity task dependent? Journal of memory and language 28(2), 127–154 (1989)

[28] Kane, M.J., Hambrick, D.Z., Tuholski, S.W., Wilhelm, O., Payne, T.W., Engle, R.W.: The generality of working memory capacity: a latent-variable approach to verbal and visuospatial memory span and reasoning. Journal of experimental psychology: General 133(2), 189 (2004)

[29] Shalizi, C.R., Crutchfield, J.P.: Computational mechanics: Pattern and prediction, structure and simplicity. Journal of statistical physics 104, 817–879 (2001)

[30] Kinney, J.B., Atwal, G.S.: Equitability, mutual information, and the maximal information coefficient. Proceedings of the National Academy of Sciences 111(9), 3354–3359 (2014)

[31] Uppal, A., Ferdinand, V., Marzen, S.: Inferring an observer’s prediction strategy in sequence learning experiments. Entropy 22(8), 896 (2020)

[32] Strelioff, C.C., Crutchfield, J.P., Hübler, A.W.: Inferring markov chains: Bayesian estimation, model comparison, entropy rate, and out-of-class modeling. Physical Review E 76(1), 011106 (2007)

[33] Strelioff, C.C., Crutchfield, J.P.: Bayesian structural inference for hidden processes. Physical Review E 89(4), 042119 (2014)

[34] Hochreiter, S., Schmidhuber, J.: Long short-term memory. Neural computation 9(8), 1735–1780 (1997)

[35] Cover, T.M.: Elements of Information Theory. John Wiley & Sons, ??? (1999)

[36] Cleeremans, A., Destrebecqz, A., Boyer, M.: Implicit learning: News from the front. Trends in cognitive sciences 2(10), 406–416 (1998)

[37] Schapiro, A., Turk-Browne, N.: Statistical learning. Brain mapping 3(1), 501–506 (2015)

[38] Perruchet, P., Pacton, S.: Implicit learning and statistical learning: One phenomenon, two approaches. Trends in cognitive sciences 10(5), 233–238 (2006)

[39] Reber, A.S.: Implicit learning of artificial grammars. Journal of verbal learning and verbal behavior 6(6), 855–863 (1967)

[40] Reber, A.S.: Implicit learning of synthetic languages: The role of instructional set. Journal of Experimental Psychology: Human Learning and Memory 2(1), 88 (1976)

[41] Cleeremans, A., McClelland, J.L.: Learning the structure of event sequences. Journal of Experimental Psychology: General 120(3), 235 (1991)

[42] Jimenez, L., Mendez, C., Cleeremans, A.: Comparing direct and indirect measures of sequence learning. Journal of Experimental Psychology: Learning, Memory, and Cognition 22(4), 948 (1996)

[43] Jiménez, L., Mendez, C.: Which attention is needed for implicit sequence learning? Journal of experimental Psychology: learning, Memory, and cognition 25(1), 236 (1999)

[44] Jiménez, L., Méndez, C.: Implicit sequence learning with competing explicit cues. The Quarterly Journal of Experimental Psychology Section A 54(2), 345–369 (2001)

[45] Visser, I., Raijmakers, M.E., Molenaar, P.C.: Characterizing sequence knowledge using online measures and hidden markov models. Memory & Cognition 35(6), 1502–1517 (2007)

[46] Frost, R., Armstrong, B.C., Christiansen, M.H.: Statistical learning research: A critical review and possible new directions. Psychological Bulletin 145(12), 1128 (2019)

[47] Mitter, S.K., Newton, N.J.: Information and entropy flow in the kalman–bucy filter. Journal of Statistical Physics 118, 145–176 (2005)

[48] Dupraz, E., Varshney, L.R.: Binary recursive estimation on noisy hardware. In: 2019 IEEE International Symposium on Information Theory (ISIT), pp. 877–881 (2019). IEEE

[49] Galesic, M., Barkoczi, D., Berdahl, A.M., Biro, D., Carbone, G., Giannoccaro, I., Goldstone, R.L., Gonzalez, C., Kandler, A., Kao, A.B., et al.: Beyond collective intelligence: Collective adaptation. Journal of the Royal Society interface 20(200), 20220736 (2023)

[50] Galesic, M., Barkoczi, D., Katsikopoulos, K.: Smaller crowds outperform larger crowds and individuals in realistic task conditions. Decision 5(1), 1 (2018)

[51] Knowlton, B.J., Squire, L.R.: Artificial grammar learning depends on implicit acquisition of both abstract and exemplar-specific information. Journal of Experimental Psychology: Learning, Memory, and Cognition 22(1), 169 (1996)

[52] Servan-Schreiber, E., Anderson, J.R.: Learning artificial grammars with competitive chunking. Journal of Experimental Psychology: Learning, Memory, and Cognition 16(4), 592 (1990)

[53] Lieberman, M.D., Chang, G.Y., Chiao, J., Bookheimer, S.Y., Knowlton, B.J.: An event-related fmri study of artificial grammar learning in a balanced chunk strength design. Journal of cognitive neuroscience 16(3), 427–438 (2004)

[54] Gobet, F., Lane, P.C., Croker, S., Cheng, P.C., Jones, G., Oliver, I., Pine, J.M.: Chunking mechanisms in human learning. Trends in cognitive sciences 5(6), 236– 243 (2001)

[55] Destrebecqz, A., Cleeremans, A.: Can sequence learning be implicit? new evidence with the process dissociation procedure. Psychonomic bulletin & review 8, 343– 350 (2001)

