## Supplemental Information for "Humans are resource-rational predictors in a sequence learning task"

### Supplementary Information

V. Ferdinand, A. Yu, and S. Marzen

October 22, 2024

#### 1 Materials and Methods

##### 1.1 Sequence generation

Participants were tasked with predicting sequences that were generated by hidden Markov models (HMMs) [17]. Such models allow for generation of stimuli that quantifiably vary in output from low complexity to high complexity, from low randomness to high randomness, and from low predictability to high predictability [4]. Furthermore, generation of these stimuli is computationally efficient and does not require the experimentalist to cleverly hand-design certain patterns: the patterns emerge from the structure of the model itself, which is easily visualized and captured in the diagrams shown in Fig. 1.

Fig. 1 shows the three HMMs that we used in this experiment: the Noisy Periodic Process (NP), the Double Process (DP) and the Even Process (EP). These models have hidden states $\sigma_t$ (here denoted by $A,\ B,\ C$) that the observer does not witness and observables $x_t$ (here denoted $0,\ 1$) that the observer sees (or receives some sort of feedback on the basis of). These are edge-emitting hidden Markov models: when in a particular hidden state $\sigma_t$ at time $t$, an observable $x_t$ is chosen according to emission probabilities $p(x_t|\sigma_t)$ and it transitions to a new hidden state $\sigma_{t+1}$ according to the transition probabilities $p(\sigma_{t+1}|\sigma_t, x_t)$. In Fig. 1, the arrows show which hidden state is transitioned to, while the labels on the arrows show the emission probability followed by the observable. For example, the emission probabilities for the Noisy Periodic Process on arrow $B \rightarrow A$ are $0.1|1, 0.9|0$ and mean that, when transitioning from state $B$ to state $A$, symbol 1 is emitted with probability $p = 0.1$ and symbol 0 is emitted with probability $p = 0.9$. Table 1 shows the relationship between the hidden state, emitted symbol, and a participant's prediction for an example sequence of length 12 generated by the Even Process. In this example, the participant makes correct predictions on all trials except $t = 3$, 5 and 10.

In the main text, we described the ideal strategy, but did not provide details on what it looked like. As an example, the prediction sequence in Table 1 is the ideal strategy for predicting the Even Process, where symbol 0 is guessed when the machine is in state $A$ and symbol 1 is guessed when the machine is in state $B$. The maximum accuracy achievable for each process (as the number of trials grows and the effects of chance decline) is 95% for NP, 81.25% for DP, and 76.92% for EP. Playing the ideal strategy requires one to deduce the hidden state of the machine and the emission probabilities of each symbol per state, however this may not be possible depending on the complexity of the HMM and the information processing limitations of the learner. When memory limitations prevent discovery of the ideal strategy, the optimality of behavior under

Table 1: Example Even Process sequence of hidden states, emitted symbols, and a participant's predicted symbols.

| trial | 1 | 2 | 3 | 4 | 5 | 6 | 7 | 8 | 9 | 10 | ... | $t$ |
|---|---|---|---|---|---|---|---|---|---|---|---|---|
| state | A | A | A | B | A | B | A | A | A | A | B | A |
| symbol | 0 | 0 | 1 | 1 | 1 | 1 | 0 | 0 | 0 | 1 | 1 | 0 |
| prediction | 0 | 0 | 0 | 1 | 0 | 1 | 0 | 0 | 0 | 0 | 1 | 0 |

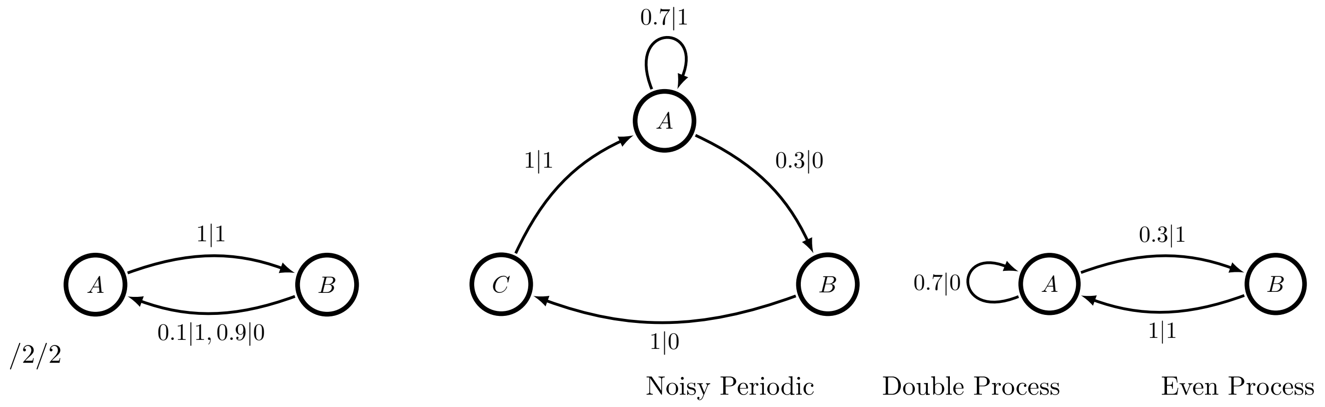


Figure 1: In each experimental condition, participants were trained on sequences generated by one of these three hidden Markov models: the Noisy Periodic Process (NP), the Double Process (DP) and the Even Process (EP).

these constraints can still be assessed by calculating the "predictive rate-accuracy curve", as described in the next section.

The three HMMs we have chosen to use in this experiment are of a particular class that allows us to easily calculate the predictive rate-accuracy curve [13]. This class is known as the $\epsilon$-Machine, probabilistic deterministic finite automaton, or unifilar hidden Markov model: given a particular hidden state $\sigma_t$ and a particular observable $x_t$, the next hidden state $\sigma_{t+1}$ is uniquely determined [19]. They are the minimal maximally-predictive model. Despite being very simple finite-state hidden Markov models, each one of the hidden Markov models used here generates an infinite-order Markov process.

For each task type, a hidden state was randomly selected, and the first observable was drawn with probabilities given by $p(x|\sigma)$. The next hidden state was transitioned to according to $p(\sigma_{t+1}|x_t, \sigma_t)$, and the process repeated 3000 times. Each participant received one unique sequence generated in this manner.

#### 1.2 Predictive rate-accuracy curves

To achieve maximal predictive accuracy, one must store "causal states", or the minimal sufficient statistics of prediction [19], which are the hidden states of the $\epsilon$-Machines in Fig. 1. However, a participant may not have enough working memory to extract the hidden states or may not use all of their working memory in the task. In this case, participants may still predict efficiently given the resources they have and attain the highest accuracy possible given the amount of memory used. Rather than simply deem the participant a worse predictor for their cognitive limitations, we ask if the way they use these limited resources is rational. More specifically, for each quantity of resources put toward the task, there will be a new (but lower) maximal predictive accuracy and participant who comes close to this new accuracy level is engaging in resource-rational prediction.

But how should we go about quantifying the amount of cognitive resources put toward the task? Ideally, we would like to know how much information the participant's brain is storing about the past of the input sequence. This is somewhat achievable in neurological studies with brain recordings [15, 12]. However, the vast majority of studies into human cognition are behavioral, where we do not have access to participants' brain state directly. If a participant's brain state is $R$, the sequence of past symbols is $\overleftarrow{X}$, the hidden state is $S$, the participant's guess is $\hat{X}$, and the true next symbol is $X$, then the Markov chain $\hat{X} \to R \to \overleftarrow{X} \to S \to X$ holds [13]. This implies through the Data Processing Inequality [3] that $I[\hat{X}; S] \leq I[R; S] \leq I[R; \overleftarrow{X}]$, or that something that looks like working memory $I[R; \overleftarrow{X}]$ is lower-bounded by something measurable in a sequence learning experiment, $I[\hat{X}; S]$, which is the mutual information between the participant's prediction and the state of the machine. (We stop coarse-graining at the causal states because at least causal states are required to predict the future [19].) The quantity $I[\hat{X};S]$ is given by $I[\hat{X};S] = \sum_{\hat{x}_t,\sigma_t} p(\hat{x}_t,\sigma_t)\log_2 \frac{p(\hat{x}_t,\sigma_t)}{p(\hat{x}_t)p(\sigma_t)}$. where $\hat{x}_t$ is the participant's prediction of which symbol will be emitted on trial $t$ and $\sigma_t$ is the state of the machine on trial $t$. Note that $I[\hat{X};\overleftarrow{X}]$ is in principle measurable too, but would be measured with much less precision since $\overleftarrow{X}$ is far larger in terms of its state space than $S$. Also, an efficient predictor as its first step knows that it only needs to pay attention to causal states and not the entire past [13].

Exactly how large their predictive accuracy can be, given a certain memory capacity $R$ (also known as a *rate*), is given by the function $A(R) = \max_{p(\hat{x}|\sigma^+):I[\hat{X};S]\leq R} \sum_{\sigma,\hat{x},x} p(\sigma)p(\hat{x}|\sigma)p(x|\sigma)\delta_{x,\hat{x}}$ where $\delta_{x,\hat{x}}$ is the Kroenecker delta function. The predictive rate-accuracy curve $A(R)$ can be computed numerically via the Blahut-Arimoto algorithm [24]. This curve delineates what accuracy levels are achievable from those that are unachievable, for any given rate $R$. All accuracies greater than $A(R)$ are forbidden, but all accuracies less than that value are allowed. Estimated combinations of rate and accuracy can lie in the forbidden region due to undersampling or nonstationarity; see Sec. 2.3. We call getting close to this predictive rate-accuracy curve "efficient prediction" because while the emphasis of getting close to the predictive rate-accuracy curve involves predicting more accurately, one can also get close to the predictive rate-accuracy curve by utilizing one's memory more efficiently [14].

Let $d(\sigma,\hat{x}) = Pr(X_{t+1} = \hat{x}|S = \sigma)$. Then $A(R)$ adopts the form of a distortion-rate curve or distortion-rate function with the said distortion measure and with $\sigma$ as input from a memoryless source and $\hat{x}$ as the decoded symbols [3]. See Sec. 2.1. As such, the Blahut-Arimoto algorithm from rate-distortion theory with a change in sign for $\beta$ to account for maximization rather than minimization can be used to find $A(R)$. The optimal $p(\hat{x}|\sigma)$ is obtained from iterating $p_{t+1}(\hat{x}|\sigma) = \frac{p_t(\hat{x})e^{\beta d(\sigma,\hat{x})}}{Z_t(\sigma)}$. $Z_t(\sigma)$ is a normalization factor.

Looking at the main text, Figure 4d-f show the predictive rate-accuracy (PRA) curve for each of the machines in Figure 1. The y-intercept of each curve is the best zero-rate strategy, which occurs when a participant allocates zero memory toward extracting the causal states from a time series and simply guesses the most common symbol on every trial. This strategy is known as *maximization* in the cognitive and decision sciences [21, 5, 6, 20, 10, 8]. The red stars in each plot show the ideal strategy, described previously, which corresponds to state-dependent maximization. Between these two extremes, the curve marks out a range of optimal strategies, where every additional bit of information put toward causal state tracking buys the learner a slightly higher predictive accuracy.

For a concrete example of an intermediate prediction strategy, consider the case of *win-stay lose-shift* (WSLS): a feedback-dependent strategy where one changes their response on $\text{trial}_{t+1}$ if it was wrong on $\text{trial}_t$ and maintains their response on $\text{trial}_{t+1}$ if it was correct on $\text{trial}_t$. A typical run from WSLS with NP looks like:

| | | | | | | | | | | | | |
|---|---|---|---|---|---|---|---|---|---|---|---|---|
| state | A | B | A | B | A | B | A | B | A | B | A | B |
| symbol | 1 | 0 | 1 | 0 | 1 | 0 | 1 | 1 | 1 | 0 | 1 | 0 |
| prediction | 0 | 1 | 0 | 1 | 0 | 1 | 0 | 1 | 1 | 1 | 0 | 1 |

where $p(\hat{x}_t|\sigma_t)$ is

| | $\hat{x}_t = 0$ | $\hat{x}_t = 1$ |
|---|---|---|
| $\sigma_t = A$ | 0.9 | 0.1 |
| $\sigma_t = B$ | 0 | 1 |

and the rate is $I[\hat{X};S] = 0.758$, meaning this strategy contains 0.758 bits of information about the hidden state of NP. In terms of accuracy, this strategy performs abysmally on NP, at 10% correct, with each lose-shift event locking the predictions into an out-of-phase alternation with the correct symbol. Of course, the performance of any given prediction strategy is dependent on the structure of the timeseries to be predicted. WSLS performs better on DP with a rate 0.95 bits and 62% accuracy, and near optimally on EP at 0.31 bits and 68% accuracy. See Figure 4d-f for the location of WSLS relative to each PRA curve.

#### 1.3 Prediction algorithms

In addition to assessing the optimality of participants' prediction behavior in a *computational* sense, as described in the previous section, we also evaluate which type of prediction *algorithm* participants' behavior is most similar to. We consider four classes of algorithms.

First is the Bayesian order-$R$ Markov model [23]. With this strategy, participants make a model of the next symbol's probability based on the last $R$ symbols seen. As described in the Supplementary Information, the prediction of the next symbol's probability is an average of the various order-$R$ Markov models. We call this strategy "n-gram", as n-gram models use the last $n$ symbols to make predictions. Second, participants might act as Bayesian Structural Inference machines [22], which means that they search through all possible $\epsilon$-Machines of up to six states and calculate the probability of each describing the observed data. We limit ourselves to six states due to the combinatorial explosion of topologies [9]. The probability of the next symbol is again found by an average over $\epsilon$-Machines. This strategy is referred to in this manuscript as "BSI". Third, participants might do logistic regression, in which the probability of the next symbol is a softmax of a linear combination of the last $k$ symbols. The parameter $k$ is a measure of memory and is chosen to maximize the log likelihood. Ridge regression was used. And finally, participants might predict like Long Short-Term Memory Units (LSTMs) [7], which are state-of-the-art recurrent neural networks that store a complicated, nonlinear memory of all past symbols in a hidden state. The output of the LSTM is then densely connected to a layer of logits. The number of nodes in the LSTM, the learning rate, the number of epochs, the number of timesteps unrolled in the LSTM during backpropagation through time were all chosen so as to ensure adequate training. LSTMs happen to be easily trainable using backpropagation through time and also have been compared to the behavior of several brain regions [18].

Ref. [25] was used to calculate log likelihoods for n-gram and logistic regression. A slight modification to the equations in Ref. [25] allowed us to calculate log likelihoods for BSI– we simply replaced the last $n$ symbols with the hidden state $\sigma$. See Supplementary Information. The n-gram and BSI models have two free parameters that were optimized using LFBGS-B as implemented by the optimization package in SciPy. Scikit-Learn was used for logistic regression with an 'L2' penalty. Tensorflow 2.0 and Keras were used to train the LSTMs, and the LSTMs were rerun five times and the best log likelihood chosen.

In more detail, we can use Ref. [25] to efficiently estimate the log likelihood of the BSI strategy in these experiments. The only difference is that the last $R$ or $n$ symbols are replaced with the causal state. To show this, we start by noting that for the BSI-average strategy (in which we average the probability of the next symbol over all possible models)

$$\log \mathcal{L} = \sum_t \log \langle P(\hat{s}_t | M_t) \rangle_{P(M_t | \overleftarrow{s}_t)}. \tag{1}$$

As in Ref. [22], we choose a Dirichlet prior over parameters $\theta$ given the model $M$ and choose a prior $M$ over models that is proportional to $e^{-\gamma|M|}$ where $|M|$ is the size of the model $M$, or the number of parameters. With these assumptions, it is straightforward to calculate

$$\langle P(\hat{s}_{t+1} | M_t) \rangle_{P(M_t | \overleftarrow{s}_t)} = \sum_M P_0(M) \frac{\alpha + \beta n(\hat{s}_{t+1} | \sigma_t)}{2\alpha + \beta n(\sigma_t)}. \tag{2}$$

The likelihood follows. Note that even though the prior is not normalized in Ref. [25], it is normalized in our code.

For all classes of algorithm considered, we assume the participant trains the model on the history of symbols emitted across all previous trials and then uses that model to predict the next trial.

Once trained, each model produces a probability distribution over the next symbol, and we assume that participants choose their prediction by weighted random sampling from that distribution. We report the "best-fit" model as the one (from this set of four) that minimizes the Akaike Information Criterion (AIC), which is $2K - 2\log L$, where $K$ is the number of parameters and $L$ is the probability of the data given the model [1]. A variant of this technique for inferring learning strategy was previously validated on simulated data [25].

#### 1.4 Working memory test

All linear mixed effects models were constructed in R [16] with the packages *lme4* [2] and *lmerTest* [11]. Three analyses were conducted: 1) differences between experimental conditions in terms of participants' distance from the curve, 2) differences between experimental conditions in terms of distance from maximum accuracy, and 3) whether rate scores are predicted by working memory scores. In analysis 1, distance from the curve on the $y$ dimension was the dependent variable, condition was the independent variable (following [?]), and random effects were entered for participant (as random intercepts). In analysis 2, the distance between participants' accuracy scores and the conditions' maximum unachievable accuracy was the dependent variable, condition was the independent variable, and random effects were entered for participant (as random intercepts). In analysis 3, participants' rate was the dependent variable, participants' working memory score was the independent variable, and random effects were entered for participant and sequence type (as random intercepts).

##### 1.5 Participants

Participants took the experiment online between 8-30 November 2021 and were recruited via Amazon's Mechanical Turk crowd sourcing platform and paid at the rate of 10 USD per hour. Participants could take the sequence learning experiment as many times as they liked and unique participant IDs were assigned using internet cookies left on the user's device. A total of 93 unique participants completed the sequence learning experiment 113 times. Of these participants, 37 opted in to take the working memory task as well. Participants were randomly assigned to a sequence condition each time they took the experiment, resulting in 44 entries for NP (35 participants), 40 entries for DP (across 34 unique participants), and 56 entries for EP (44 participants).

#### 2 Near-optimal efficient prediction

##### 2.1 Connection to rate-distortion theory

In rate-distortion theory, a memoryless source $X$ spits out $n$ symbols $x_1, ..., x_n$ and sends these symbols through a channel. The channel produces one of $M$ words and is said to have a rate of $\log M/n$ bits per input symbol. This word is then decoded by a decoder as $\hat{x}_1, ..., \hat{x}_n$. The decoder is said to have a distortion of $\frac{1}{n}\sum_{i=1}^n d(x_i, \hat{x}_i)$ for some distortion measure $d : \mathcal{X} \times \mathcal{X} \to \Re$. We define the rate-distortion function or rate-distortion curve to be the minimal rate achievable given a particular distortion,

$$R(D) = \min_{\text{channel,decoder:distortion}\leq D} \text{rate}. \tag{3}$$

Likewise, one can define the distortion-rate function or distortion-rate curve as the inverse,

$$D(R) = \min_{\text{channel,decoder:rate}\leq R} \text{distortion}. \tag{4}$$

The problem of finding $R(D)$ or $D(R)$, which separates achievable from unachievable combinations of channels and decoders, looks to be intractable. How could you ever search over all possible channels and decoders? But the rate-distortion theorem turns this seemingly intractable problem into one that can be computed using the Blahut-Arimoto algorithm:

$$R(D) = \min_{p(\hat{x}|x):\mathbb{E}[d(x,\hat{x})]\leq D} I[X;\hat{X}]. \tag{5}$$

The distortion-rate function or distortion-rate curve is then the inverse of this. Note that the rate-distortion theorem breaks down if the source is memoryful (as it is for us) or if the blocklength $n$ is not infinite. If $n$ is finite, then $R(D)$ is replaced by $R_n(D)$, which approaches $R(D)$ as $n \to \infty$ from above. If the source is memoryful, then one replaces $I[X;\hat{X}]$ with $\lim_{n\to\infty} \frac{I[X_{1:n};\hat{X}_{1:n}]}{n}$. Further, and perhaps most importantly, we are working with predictive distortions in which the distortion measure depends on $p(\hat{x}|x)$. As such, we are incapable of using the rate-distortion theorem to justify using our predictive rate-accuracy curve.

Instead, we merely assert that there is an interesting information bottleneck that justifies using rate (now historically named) to measure the size of the bottleneck. Pasts $\overleftarrow{X}_t$ of the input string convey information

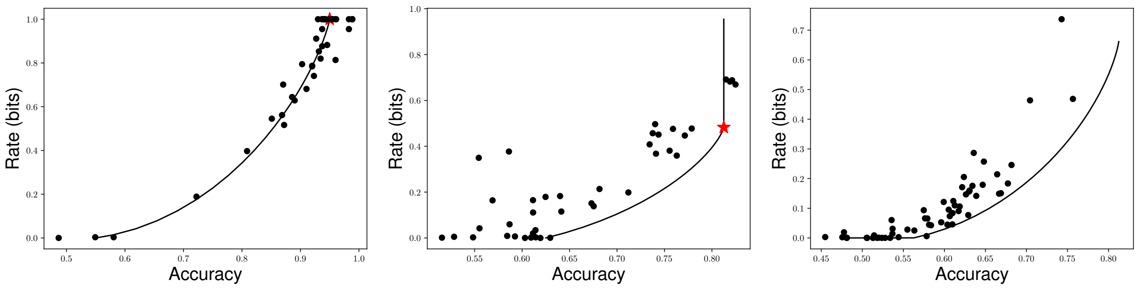


Figure 2: At left, Noisy Periodic. In the middle, the Double Process. And at right, the Even Process. These are simply the plots in Fig. 2 of the main text with $x$ and $y$ axes transposed.

about the future $X_{t+1}$ only through the hidden/causal state $S_t$. The brain state $R_t$ only can see the past, and the prediction $\hat{X}_t$ is a function only of the brain state. Note that the $x$ in the previous paragraph is now $\sigma$, the causal state, and $\hat{x}$ is the prediction still. We have the Markov chain $\hat{X}_t \to R_t \to \overleftarrow{X}_t \to S_t \to X_{t+1}$, which gives the inequality in the main text. In words, information about the future input can only be obtained from information about the hidden/causal state, and this information is squeezed through a bottleneck which is the prediction itself. The size of the bottleneck is approximated by $r = I[\hat{X}_t; S_t]$. Predictive accuracy $a$ is a reasonable measure of how well someone is predicting.

#### 2.2 The predictive rate-accuracy curve's complement and distances from the curve

In the main text, we showed the predictive rate-accuracy curve, with accuracy on the $y$-axis and rate on the $x$-axis. But it is more standard in information theory to show the rate-distortion curve, so we show the corresponding plots below in Fig. 2.

These plots allow us to visualize the normalized distance for the rate, just as the figures in the main text allow us to visualize the normalized predictive accuracy. The idea behind normalized predictive accuracy distances is that there is some minimum accuracy (0) and some maximum accuracy for a given rate, $A(r)$; we compute how far of the way the actual accuracy is up to the maximum possible accuracy, so $\frac{A(r)-a}{A(r)}$. The idea behind normalized rate distance is that there is some minimum rate for a given accuracy $R(a)$ and some maximum possible rate $R_{max} = \max_{p(x|\sigma)} I[X; S]$; we compute how far of the way towards the minimal rate the actual rate is, so $\frac{r-R(a)}{R_{max}-R(a)}$. Violin plots of the normalized predictive accuracy distances and normalized rate distances are shown in Fig. 3. They are signed, and due to undersampling and nonstationarity, might be negative.

#### 2.3 Nonstationarity

A number of rate and accuracy combinations reported in the main text are in the forbidden region. While this might be due to undersampling, this is more likely due to nonstationarity in guessing strategy.

There was notable nonstationarity in many of the participant's guessing strategies. See Fig. 4 for estimated $\hat{p}(0|\sigma)$ based on the number of times 0 followed hidden state $\sigma$ in trial lengths of 100 as a function of the starting trial number. We also calculate a nonstationarity score, the expected variance of $p(0|\sigma)$ over the last half of the experiment and show the histogram of these stationarity scores in violin plots.

As can be seen from the last subfigure in Fig. 4, nonstationarity was observed for the Double Process and Even Process, and less so for the Noisy Periodic process. The effects of nonstationarity on predictive rate-accuracy curves can be enormous. That is why computation of Fig. 4 is necessary to show that some degree of stationarity was observed.

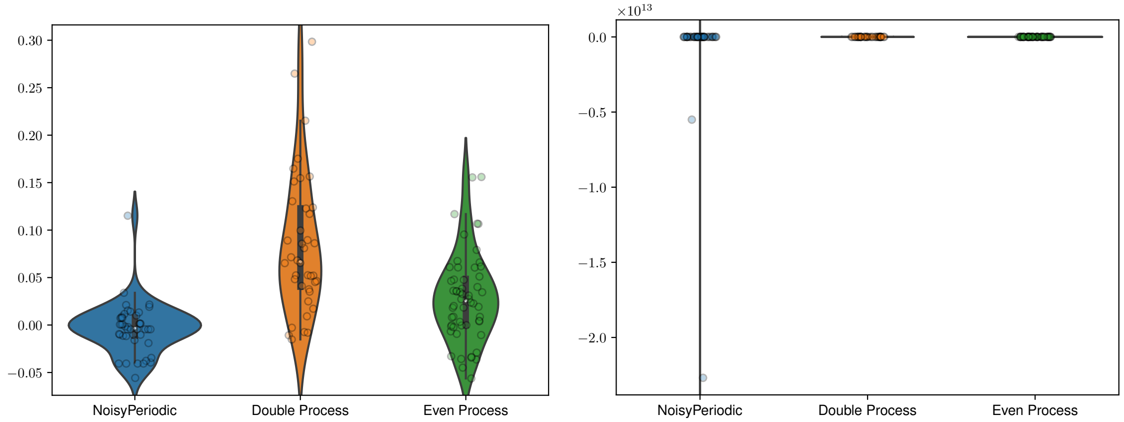


Figure 3: At left, violin plots of $(A(r) - a)/A(r)$. At right, violin plots of $(r - R(a))/(R_{max} - R(a))$.

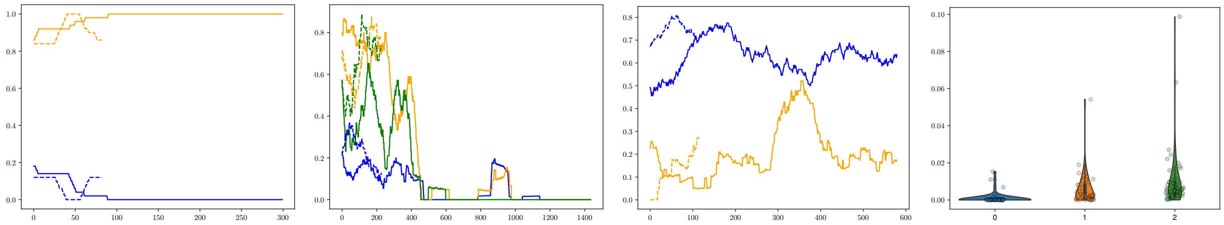

Figure 4: We describe from left to right. First, Noisy Periodic's $\hat{p}(0|\sigma)$, with $\sigma = A$ in blue and $\sigma = B$ in orange. Then, the Double Process' $\hat{p}(0|\sigma)$, with $\sigma = A$ in blue, $\sigma = B$ in orange, and $\sigma = C$ in green. Third, the Even Process' $\hat{p}(0|\sigma)$, with $\sigma = A$ in blue and $\sigma = B$ in orange. In all plots, the solid line shows a participant whose guessing strategy becomes essentially stationary by the end of the experiment and the dotted line shows a participant whose strategy is still changing markedly by the end of the experiment. Finally, a violin plot of the nonstationarity scores $(\mathbb{E}[\mathrm{Var}[p(0|\sigma)]])$ for each task, showing that the Noisy Periodic process leads to far more stationary guessing strategies than the Double or Even Process.

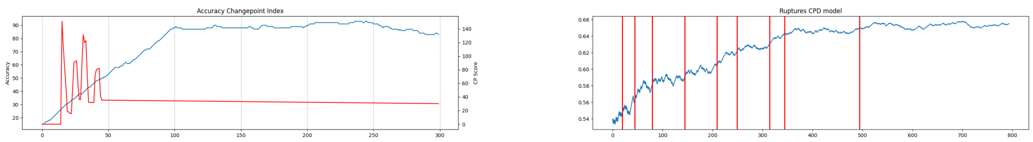


Figure 5: At left, SDAR on the rolling predictive accuracies for one participant. Change points appear to be incorrectly identified. At right, Ruptures on the rolling predictive accuracies for another participant. Ruptures tended to be overaggressive in identifying change points.

To see the effects of nonstationarity theoretically, suppose that the participant in question has a nonstationary guessing strategy, $p_t(x|\sigma)$. The average guessing strategy is $\langle p_t(\hat{x}|\sigma)\rangle_t$, and this is what was used to make Fig. 2 of the main text. Rates were computed from this conditional probability distribution, and accuracies were computed from comparing $\hat{x}_t$ to $x_t$. If $p_t(\hat{x}|\sigma)$ changes quickly enough, then just by chance, it is possible to exceed the accuracy limits set by the predictive rate-accuracy curve, which assumes a stationary strategy, *even though* any particular strategy $p_t(\hat{x}|\sigma)$ is subject to the limits set by the predictive rate-accuracy curve.

Suppose that the last half is divided into five parts which each have a stationary strategy. Then, the rate in Fig. 2 of the main text would underreport the true rate. The accuracy for the average strategy is essentially the average of the accuracies over all trials $t$:

$$A = \sum_{\hat{x},\sigma} p(\sigma)\langle p_t(\hat{x}|\sigma)\rangle_t Pr(X_{t+1} = \hat{x}|S_t = \sigma) = \langle \sum_{\hat{x},\sigma} p(\sigma) p_t(\hat{x}|\sigma) Pr(X_{t+1} = \hat{x}|S_t = \sigma)\rangle_t = \langle A_t \rangle_t. \tag{6}$$

However, the rate of the average strategy is not the average of the rates over all trials $t$ typically, and in fact, according to Jensen's inequality and the convexity of the mutual information $I[X;S]$ in $p(x|\sigma)$, we have

$$\langle I[p_t(x|\sigma)]\rangle_t \geq I[\langle p_t(x|\sigma)\rangle_t]. \tag{7}$$

As a result, the rate in Fig. 2 of the main text would be less than the means of the actual rates, while the accuracy would be spot on.

Due to the difficulties with interpreting the predictive rate-accuracy plots when there is nonstationarity, we tried to limit the trials to those that appeared to have a stationary guessing strategy. First, we tried to implement popular change-point detection algorithms on rolling predictive accuracy scores as implemented in sci-kit learn. Both SDAR and Ruptures tended to either miss or find too many change points that did not correspond to change points that one would see by eye. See Fig. 5. Further, we tried to use the bonus round only, but this eliminated many data points and still left data points that had significant nonstationarity.

### References

[1] D Anderson and K Burnham. Model selection and multi-model inference. *Second. NY: Springer-Verlag*, 63(2020):10, 2004.

[2] Douglas Bates, Martin Mächler, Ben Bolker, and Steve Walker. Fitting linear mixed-effects models using lme4. *Journal of Statistical Software*, 67(1):1–48, 2015.

[3] Thomas M Cover. *Elements of information theory*. John Wiley & Sons, 1999.

[4] David P Feldman, Carl S McTague, and James P Crutchfield. The organization of intrinsic computation: Complexity-entropy diagrams and the diversity of natural information processing. *Chaos: An Interdisciplinary Journal of Nonlinear Science*, 18(4), 2008.

[5] R Allen Gardner. Probability-learning with two and three choices. *The American Journal of Psychology*, 70(2):174–185, 1957.

[6] Richard J Herrnstein. Rational choice theory: Necessary but not sufficient. *American Psychologist*, 45(3):356, 1990.

[7] Sepp Hochreiter and Jürgen Schmidhuber. Long short-term memory. *Neural computation*, 9(8):1735–1780, 1997.

[8] Carla L Hudson Kam and Elissa L Newport. Regularizing unpredictable variation: The roles of adult and child learners in language formation and change. *Language learning and development*, 1(2):151–195, 2005.

[9] Benjamin D Johnson, James P Crutchfield, Christopher J Ellison, and Carl S McTague. Enumerating finitary processes. *arXiv preprint arXiv:1011.0036*, 2010.

[10] Daniel Kahneman. Maps of bounded rationality: Psychology for behavioral economics. *American economic review*, 93(5):1449–1475, 2003.

[11] Alexandra Kuznetsova, Per B. Brockhoff, and Rune H. B. Christensen. lmerTest package: Tests in linear mixed effects models. *Journal of Statistical Software*, 82(13):1–26, 2017.

[12] Martina Lamberti, Shiven Tripathi, Michel JAM van Putten, Sarah Marzen, and Joost le Feber. Prediction in cultured cortical neural networks. *PNAS nexus*, 2(6):pgad188, 2023.

[13] Sarah E Marzen and James P Crutchfield. Predictive rate-distortion for infinite-order markov processes. *Journal of Statistical Physics*, 163:1312–1338, 2016.

[14] Sarah E Marzen and James P Crutchfield. Probabilistic deterministic finite automata and recurrent networks, revisited. *Entropy*, 24(1):90, 2022.

[15] Stephanie E Palmer, Olivier Marre, Michael J Berry, and William Bialek. Predictive information in a sensory population. *Proceedings of the National Academy of Sciences*, 112(22):6908–6913, 2015.

[16] R Core Team. *R: A Language and Environment for Statistical Computing*. R Foundation for Statistical Computing, Vienna, Austria, 2018.

[17] Lawrence R Rabiner. A tutorial on hidden markov models and selected applications in speech recognition. *Proceedings of the IEEE*, 77(2):257–286, 1989.

[18] Kanaka Rajan, Christopher D Harvey, and David W Tank. Recurrent network models of sequence generation and memory. *Neuron*, 90(1):128–142, 2016.

[19] Cosma Rohilla Shalizi and James P Crutchfield. Computational mechanics: Pattern and prediction, structure and simplicity. *Journal of statistical physics*, 104:817–879, 2001.

[20] David R Shanks, Richard J Tunney, and John D McCarthy. A re-examination of probability matching and rational choice. *Journal of Behavioral Decision Making*, 15(3):233–250, 2002.

[21] Herbert A Simon. A behavioral model of rational choice. *The quarterly journal of economics*, pages 99–118, 1955.

[22] Christopher C Strelioff and James P Crutchfield. Bayesian structural inference for hidden processes. *Physical Review E*, 89(4):042119, 2014.

[23] Christopher C Strelioff, James P Crutchfield, and Alfred W Hübler. Inferring markov chains: Bayesian estimation, model comparison, entropy rate, and out-of-class modeling. *Physical Review E*, 76(1):011106, 2007.

[24] Naftali Tishby, Fernando C Pereira, and William Bialek. The information bottleneck method. *arXiv preprint physics/0004057*, 2000.

[25] Abhinuv Uppal, Vanessa Ferdinand, and Sarah Marzen. Inferring an observer's prediction strategy in sequence learning experiments. *Entropy*, 22(8):896, 2020.
